# Distinct growth stages shaped by an interplay of deterministic and neutral processes are indispensable for functional anammox biofilms

**DOI:** 10.1101/2020.06.03.131896

**Authors:** Robert Niederdorfer, Lisa Fragner, Ling Yuan, Damian Hausherr, Jing Wei, Paul Magyar, Adriano Joss, Feng Ju, Helmut Bürgmann

**Affiliations:** Eawag, Swiss Federal Institute for Aquatic Science and Technology, Department of Surface Waters-Research and Management, 6047 Kastanienbaum, Switzerland; Environmental Microbiome and Biotechnology Laboratory (EMBLab), School of Engineering, Westlake University, 18 Shilongshan Road, Hangzhou 310024, China; Eawag, Swiss Federal Institute for Aquatic Science and Technology, Department of Process Engineering, 8600 Dübendorf, Switzerland; Empa, Swiss Federal Laboratories for Materials Science and Technology, Laboratory for Air Pollution & Environmental Technology, 8600 Dübendorf, Switzerland; Department of Environmental Sciences, University of Basel, Basel, Switzerland

**Author notes:** Joint first authorship (Robert Niederdorfer, Lisa Fragner).

## Abstract

Complex microbial biofilms orchestrating mainstream anaerobic ammonium oxidation (anammox) represent one of the most promising energy-efficient mechanisms of fixed nitrogen elimination from anthropogenic waste waters. However, little is known about the ecological processes that are driving microbial community assembly leading to functional anammox biofilms in engineered ecosystems. Here, we use fluorescence in situ hybridization and 16S rRNA sequencing combined with network modelling to elucidate the contribution of stochastic and deterministic processes during anammox biofilm development from first colonization to maturation in a carrier-based anammox reactor. We find that distinct stages of biofilm development emerge naturally in terms of structure and community composition. These stages are characterized by dynamic succession and an interplay of stochastic and deterministic processes. The staged process of biofilm establishment appears to be the prerequisite for the anticipated growth of anammox bacteria and for reaching a biofilm community structure with the desired metabolic capacities. We discuss the relevance of this improved understanding of anammox community ecology and biofilm development concerning its practical application in the start-up and configuration of anammox biofilm reactors.

## Introduction

Anaerobic ammonium oxidation (anammox) involves the simultaneous oxidation of ammonium (NH_4_^+^) and reduction of nitrite (NO_2_^−^) under oxygen-limiting conditions and is orchestrated by a unique lineage of bacteria (AMX), which all belong to a monophyletic group within the *Planctomycetes* (Kartal et al. 2011; Strous et al. 2006). Used under mainstream conditions in waste water treatment plants (WWTP), anammox is also one of the most promising energy-efficient mechanisms of fixed nitrogen elimination in engineered systems and represents a step towards energy autarky in the treatment of municipal waste waters (Siegrist et al. 2008; Lotti et al. 2014). AMX are generally characterized by a lack of pure cultures, very slow growth rates and low cell yields (Kuenen 2008), which makes sufficient retention of biomass one of the main challenges regarding their engineered application (Abma et al. 2007). In response, engineers facilitate the formation of complex microbial communities and exploit the ecological growth strategy of AMX bacteria to grow in matrix-enclosed biofilms on artificial carrier material or as suspended granular sludge, to increase biomass retention (Strous et al. 1997; Flemming et al. 2016). In addition to the improved retention time of biofilms, lower space requirements, lower sludge production and the resilience to changes in the reactor configuration are only some of the benefits of biofilm-based treatment systems (Wilderer and McSwain 2004; Chen and Chen 2000; Yingxin Zhao et al. 2019).

The microbial communities that form the AMX biofilms in wastewater treatment systems are very diverse, with a majority of heterotrophic bacteria that are crucial for the formation of the biofilm, and therefore creating the habitat for AMX, but do not contribute directly to the anammox process itself (Laureni et al. 2015; Mozumder et al. 2014; Ni, Ruscalleda, and Smets 2012).

While there are numerous studies focusing on microbial community assembly in suspended systems (Chu et al. 2015; Luo et al. 2017; Gonzalez-Martinez et al. 2015), the ecological processes underlying AMX biofilm formation on synthetic carrier material in engineering applications are still poorly understood. The rich diversity, potential for numerous synergetic and antagonistic interactions (Battin et al. 2016; Flemming and Wuertz 2019; Faust et al. 2018) and stochastic effects (Sloan et al. 2006; Woodcock and Sloan 2017) that govern community assembly complicate efforts to fully understand AMX biofilm formation for engineered applications (Lawson et al. 2019). A better mechanistic understanding of these factors is needed, also with respect to the growing interest in microbial community engineering for waste water treatment (Moralejo-Gárate et al. 2011; Kleerebezem and van Loosdrecht 2007; Vlaeminck, De Clippeleir, and Verstraete 2012).

A spatial, i.e. microscopy-based investigation of AMX bacterial biofilm provides important information about the structural and spatial requirements of AMX within such a complex community (Almstrand et al. 2013; Kindaichi et al. 2007; Laureni et al. 2015). Together with insights about temporal community dynamics, derived from sequencing data, it is possible to draw conclusions about the ecological processes that lead to an established biofilm community (Niederdorfer, Peter, and Battin 2016; Shade et al. 2014; Brislawn et al. 2019; Datta et al. 2016). Within this methodological framework, co-occurrence network analyses combined with fundamentals of community succession ecology have proven to be a powerful tool to elucidate the mechanistic background of ecological interactions between microbial community members in even more detail (Ju and Zhang 2015; Röttjers and Faust 2018; Faust and Raes 2012; Widder et al. 2016). Therefore, both spatial and compositional data is required to gain an in-depth understanding of biofilm formation.

In order to understand the ecological processes that lead to anticipated AMX growth in biofilms, we investigated the microbial community assembly on synthetic biofilm carriers from a pilot scale mainstream anammox reactor, over the course of a year from first colonization to mature biofilms. We hypothesized i) that heterotrophic bacteria are the pioneering colonizers that initiate biofilm formation and create the required base layer for second wave colonizers including AMX, and ii) that structure and succession of biofilms can be divided into three distinct growth stages - colonization, succession and maturation. We expected colonization to be driven by initial colonizers capable to attach and grow on the industrial carrier surface. Succession was expected to be driven by increasingly intense competition for space and nutrients in the juvenile biofilm. The mature phase represents the convergence to a well-adapted quasi-equilibrium state determined by the environmental factors in the reactor. We further assumed that both stochastic (e.g., birth, growth, death and immigration) and deterministic processes (e.g., environmental factors, biotic interactions, niche differentiation, historical (e.g., priority) effects) are important in varying degrees for community assembly across the different stages. In accordance with hypothesis ii we therefore further hypothesized iii) that stochastic and deterministic processes are changing in their relative contribution on the community assembly over the different stages of biofilm development.

By combining fluorescence *in situ* hybridization and 16S rRNA sequencing of the developing microbial community, we aimed to determine the spatial and temporal succession dynamics of AMX biofilms on a synthetic carrier material. We are dissecting the complexity of the microbial community into subpopulations based on their occurrence patterns and calculated their contribution to the total community turnover over the course of biofilm growth. Using neutral, co-occurrence and checkerboard score modeling of AMX biofilm community assembly allowed us to disentangle deterministic and stochastic ecological factors that paved the way for a mature biofilm community. To our knowledge, this is one of the first studies using a toolbox of various community modelling approaches combined with spatial analysis to unravel assembly mechanisms of surface-attached AMX biofilms in engineered ecosystems. We believe our results provide valuable scientific information for future engineering studies in terms of anammox reactor configuration and reactor start up. Furthermore, it highlights the importance of bioreactor systems for detailed microbiome studies.

## Material and Methods

### Experimental Set-Up and Sampling

The experiment was conducted in an 8 m^3^ mainstream anammox pilot reactor, which is part of the pilot wastewater treatment plant at Eawag in Dübendorf. A separate pilot scale (8m^3^) high rate activated sludge reactor, fed with fresh mainstream municipal wastewater from the municipality Dübendorf, provided the chemical oxygen demand-depleted influent for the anammox reactor. The AMX reactor is therefore exposed to mainstream-like conditions, including the seasonal temperature variations of the in-flowing wastewater (~25-12°C). The reactor is operated in sequencing batch reactor (SBR) mode and is discontinuously filled and emptied every 6-7 hours, depending on the process cycle length. Each SBR cycle was controlled by an automated control sequence that consisted of five steps: (1) settling (60 min), (2) effluent discharge, (3) feeding, (4) reaction phase and (5) mixing. To prevent washout of the biomass during waste water decanting, 150 m^2^/m^3^ of FLUOPUR^®^ carrier material (WABAG Water Technology Ltd., Switzerland) fleece tiles made of synthetic polymeric fibers with a size of 12×12×0.1 mm are used in the anammox reactor. At the start of our experiment, the AMX reactor had been operating successfully for over 2 years (~100gNH4-N m^3^ d^−1^).

For the colonization experiment, we added 35,000 marked (cut corners) blank biofilm carriers to the system and sampled over a period of a year. Carrier sampling (n=6) was performed twice per week over the first eight weeks and on a weekly basis thereafter for one year. Old carriers that had remained in the tank since June 2016 were also sampled at each time point for comparison. The samples were immediately transferred to tubes, snap frozen in liquid nitrogen and stored at −80 °C for further analysis.

### Sample fixation and cryosectioning

Six sampling points (one, three, six, nine, twelve and 24+ months) were chosen for cryosectioning and analysis by fluorescence in-situ hybridization (FISH). Duplicate carriers were analyzed for each time point, with an additional blank carrier as a control. The carriers were cut into three sections, with the middle section being used for fixation, cryosectioning and FISH-Confocal laser scanning microscopy (CLSM). The two remaining carrier sections were stored at –80 °C for subsequent DNA extraction.

The carrier sections were fixed in a 4 % formaldehyde solution for three hours at room temperature. The samples were then washed by rinsing three times with a 1x phosphate-buffered saline (PBS) solution and stored at 4 °C in a 1:1 mixture of 1x PBS solution and 96 % ethanol. Samples were then embedded in an optimal cutting temperature embedding matrix (Cell Path Ltd., United Kingdom) and frozen at –10 °C.

Cryosectioning was performed on a Thermo Scientific™ CryoStar™ NX70 Cryostat (Thermo Fisher Scientific Inc., United States) at the ScopeM facility (ETH Zurich). Afterwards, the coated and dehydrated slides were stored at −20 °C for later microscopy.

FISH was performed according to established standard operational procedures (Nielsen 2009) using the oligonucleotide probes listed in Supplementary Table 1 to identify all bacteria (EUB), nitrite-oxidizing bacteria (NOB) and AMX in the cryosectioned samples (Laureni et al. 2015). In addition, a negative control for each sample without the addition of FISH-probes was performed.

### Microscopy and Image Analysis

CLSM of the FISH-slides was performed with a Leica SP5 DMI 6000 (Leica Microsystems GmbH, Germany) and the software LAS AF v2.7.9. Images with a resolution of 1024×1024 pixels (4.096 pixels/μm) were collected using a 63x/1.40 oil objective. Z-stacks were taken at five randomly selected locations on each of three sample slices of each examined carrier, resulting in 15 recorded z-stacks per sample and 30 per time point. Z-stack size varied with thickness of the cryosections between 20-30 μm. A z-step size of 0.5 μm was chosen, resulting in z-step numbers between 40 and 60. The image files were visualized and extracted in Fiji; a distribution of the open-source software ImageJ (Schindelin et al. 2012). Daime (version 2.1) (Daims, Lücker, and Wagner 2016) was used for 3D visualization and digital image analysis of the z-stacks. Prior to volume measurement, the z-stacks of each channel (Cy3, Cy5, FITC) and each sample were prepared. Initially it was necessary to reduce the resolution of each image to 512×512 pixels, while the 3D median filter was used to reduce noise. 3D segmentation with edge detection mode was used to distinguish between biomass and background, with objects up to 50 voxels being considered as background artefacts, and therefore ignored. The detected objects within the segmented z-stacks were controlled visually and any remaining wrong signals from cut carrier fibers and/or air bubbles were identified and rejected manually. Thereafter the total volume of the objects from each channel was determined and the mean volume of all 30 random images per time point was calculated to estimate the development of biomass volume over time taken the maximum reachable volume from the old carrier. We calculated the ratio between the mean biomass volume (mm^3^) and number of objects for every time point to investigate their correlation.

### Biomass sampling, extraction and sequencing

Nucleic acids were extracted based on a method modified from Griffiths et al. (2000). Duplicate biofilm carriers (n = 3) from every time point were cut into small pieces on a liquid nitrogen bath. Carrier pieces were transferred to 1.5 ml Matrix E lysis tubes (MPbio) and 0.5 ml of both hexadecyltrimethylammonium bromide buffer and phenol:chloroform:isoamylalcohol (25:24:1, pH 6.8) was added. Cells were lysed in a FastPrep machine (MPbio), followed by nucleic acid precipitation with PEG 6000 on ice. Nucleic acid pellets were washed three times with ethanol (70%) and dissolved in 50 μl DEPC treated RNAse free water. DNA quality and quantity was assessed by using agarose gel electrophoresis and a Nanodrop ND-2000c (Thermo Fisher Scientific, USA). 16S rRNA gene amplicon sequencing was performed by Novogene Ltd. (Hong Kong) on the Illumina NextSeq platform, based on a pair-end algorithm (250bp, V3-V4).

### Sequence analysis and ecostatistics

Raw sequences were analyzed within the QIIME2 framework (Bolyen et al. 2019). Primer sequences were removed with the cutadapt QIIME2 plugin. After demultiplexing, read pairs were joined, low-quality reads were filtered out and all high quality reads were analyzed with the DEBLUR software to produce amplicon sequence variants (ASVs) based on Illumina Miseq/Hiseq error profiles. Taxonomical assignment of the ASVs was performed within QIIME2 environment with the SILVA 16S V3/V4 classifier (accessed January 2020). After filtering of unclassified and contaminated ASVs, the resulting sequence table consisted of 1038 ASVs. All subsequent analyses were performed on the relative abundances sequence table (rarefied to 15595 reads) to ensure comparability between the samples. Based on the relative abundances we plotted the top bacterial phyla over time for each sample. Non-metric multidimensional scaling (nMDS) analysis on Bray–Curtis dissimilarity matrices served to visualize patterns of community composition, and PERMANOVA tested for differences among communities. Based on the results of nMDS and visual observations we categorized the dataset into colonization (Col), succession (Suc), maturation (Mat). Old carriers were treated as a separate category (Old). We calculated the coefficient of variation in abundance (CV) for all ASVs within the development phases Col, Suc, Mat and Old) to assess their variation during biofilm development. We calculated the relative proportions for every ASVs between the different growth stages and visualized them in a ternary plot.

### Residents, transient and exclusive ASVs

Dependent on the occurrence of ASVs within colonization, succession, maturation, and old samples we organized them into residents (all ASVs that were present at least in 80% of all samples over all development phases), transients (ASVs that were present in at least 80% of the samples of any two phases and absent during the other) and exclusives (ASVs that occurred in >80% of samples of only one biofilm development phase and were absent during the others). We calculated the Bray–Curtis dissimilarity between adjacent time points and calculated the fraction of dissimilarity contributed by ASVs identified as residents, transients, exclusives and the ones that could not be categorized (other). To clarify the distribution we divided the transients into early transients (only occurring during Col and Suc) and late transients (only occurring during between Suc and Mat).

### Neutral modeling on the stochasticity of the community assembly process

To assess the relative importance of stochasticity and determinism for anammox community assembly, we firstly evaluated the fit of a Sloan neutral community model (Sloan et al. 2006) for each stage. The neutral model supposes community assembly is determined by dispersal and drift, i.e. two stochastic components of ecological processes. Analysis was performed in R using a published script (Burns et al. 2016). Output plots mainly show: (i) the fit of neutral model, r2, and the migration rate, m; (ii) neutral model predictions and its corresponding 95% confidence intervals (lines) and actual distributions (points) for each successional stage. We then calculated an index normalized stochasticity ratio (NST) for each stage to verify the results of Sloan neutral model. NST is a normalized index that ranges from 0 to 1 which represent the proportion of ecological stochasticity quantitatively with great accuracy and precision. Analysis was conducted in R using the package NST (Ning et al. 2019).

### Co-occurrence network and C-score modeling on process determinism

To construct co-occurrence networks for microbial community in the anammox biofilm system, Spearman’s rank coefficients (ρ) between all ASVs abundances with occurrence in more than 50% of samples were calculated. Correlations with ρ>0.8 and P-value<0.05 were considered statistically significant and robust. A series of topological properties including MD (modularity), CC (average clustering coefficient), APL (average shortest path length) and ND (network density) were calculated and compared across each stage. Statistical analyses were conducted in R using package VEGAN (Oksanen et al. 2013), igraph (Csardi and Nepusz 2006) and Hmisc (Harrell and Dupont 2016). Networks were visualized and modularized in Gephi 0.92 (Bastian, Heymann, and Jacomy 2009). Moreover, a modified python script from (Ju and Zhang 2015) was used to calculate the random (R%) and observed (O%) incidences of intra-group and inter-group co-occurrence patterns between microbial entities (i.e., ASVs). The degree of agreement between O and R (O/R ratio) allows for checking non-random assembly patterns in complex communities (Ju and Zhang 2015).

The checkerboard score (C-score) counts the number of checkerboard units (i.e., 2 × 2 matrix) in a community. It was used as the metric of species segregation (Stone and Roberts 1990), and its variance (C-scorevar) was used as the metric of both species segregation and aggregation in the community. The C-score and Cvar-score tests were performed using R package ‘EcoSimR’ (http://ecosimr.org/) with constant row and column sums, sequential swap randomization algorithm, and a burn-in of 100,000 swaps. The higher the observed standardized effect size (SES) values of the C-score the greater the degrees of species segregation than expected by chance. Higher SES values of C_var_-score indicate greater degrees of both species segregation and aggregation. Python and R scripts used for co-occurrence network and C-score analysis are freely available at https://github.com/RichieJu520.

## Results

### Temporal development of anammox biofilm structure revealed by FISH-CLSM

During the first month of biofilm growth pioneering bacteria attached to the carrier material and accumulated into small microcolonies in the voids between the fibers (Figure 1). While Anammox bacteria did not contribute to the early phase of biofilm development, NOB were already part of the community in small patches (Figure 1). After three months anammox consortia also started to appear as microcolonies, while existing bacterial patches grew further in size. After six months, larger and sporadically connected agglomerations of EUB had developed along with the formation of numerous small AMX and NOB colonies attached to or enclosed within the EUB patches. After nine months of biofilm growth, AMX clusters displayed a significant increase in volume. EUB also increased in biomass by combining patches to form bigger agglomerations and by building new colonies, while the NOB fraction continued to form additional small colonies. After twelve months, NOB patches appeared to start connecting to form larger agglomerations; however, not comparable to the extent of observations for AMX and EUB agglomerations. Finally, in the old biofilms (24+) AMX had formed additional small patches while the larger AMX agglomerations and the EUB agglomerations grew extensively more in size, taking over more than 50% of the carrier volume.

**Figure 1.**
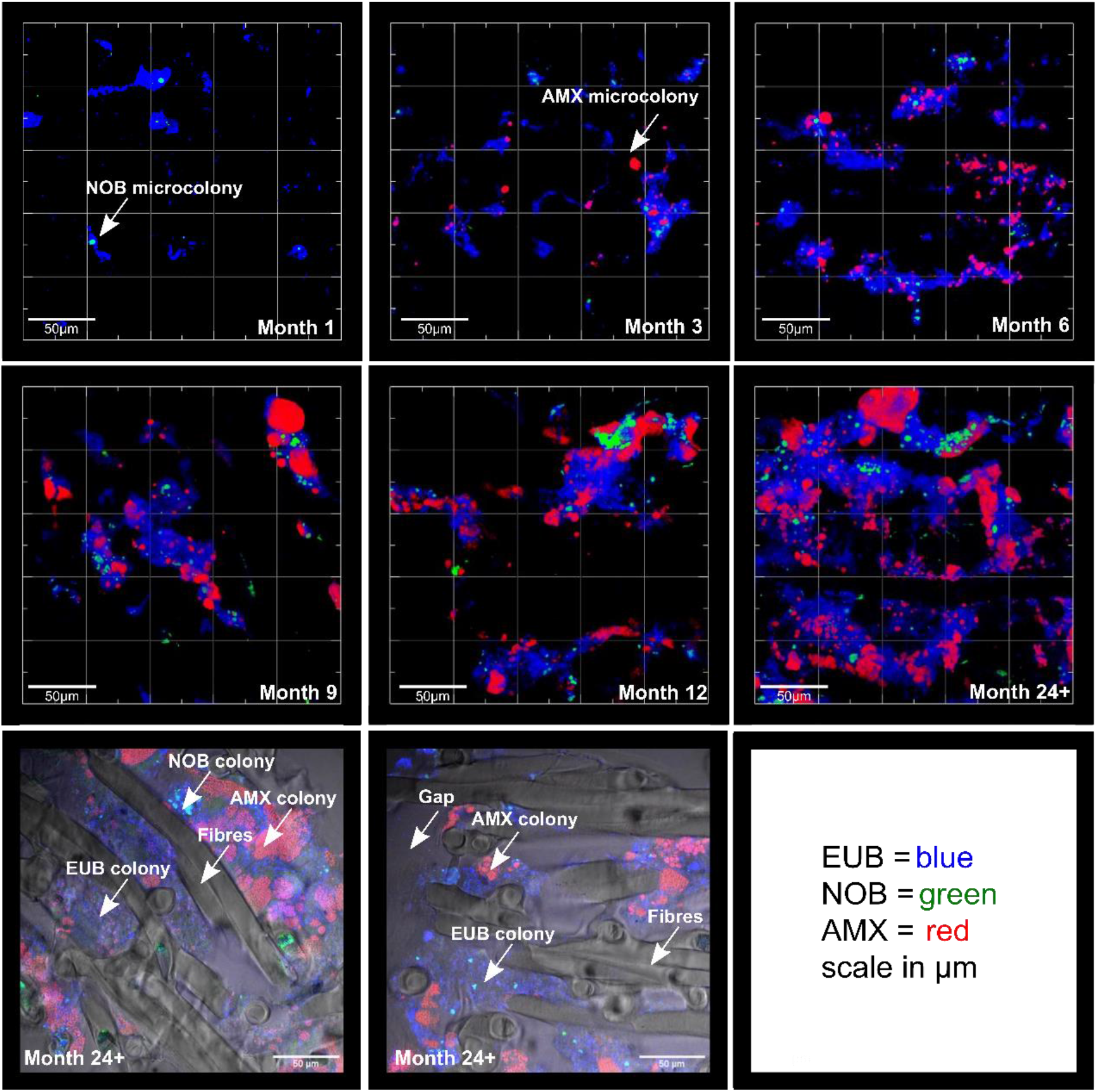
**Row 1 and 2**: Representative FISH-CLSM images of biofilm development taken place over the course of a year on synthetic carrier material. Colours denote stained bacterial fraction in the community: EUB = blue, NOB = green, AMX = red. The scale unit is μm. **Row 3:** Brightfield captions of the old carriers overlaid with FISH-CLSM pictures. Arrows highlight certain regions on the carrier.

Comparison of bright field and FISH microscopy showed that bacteria only grew around and not within the fibers of the carrier, which led to the observed gaps between the bacterial accumulations (Figure 1, Month 24+). The available pore space in the carrier appeared to be nearly filled with bacterial biomass in the mature and old biofilm.

We used image analysis to determine the mean total volume and number of detected objects for each time point (Table 1). The average volume of objects (colonies or patches) expressed as the ratio between average volume and average number of detected objects reflected the qualitative visual observations (Figure 2A). EUB initiated the biofilm formation but expanded slowly in volume (Figure 2B). Between three and six months, the ratio increased for all three bacterial consortia. The NOB volume/object ratio remained far lower than for EUB and AMX and was relatively stable over time although the overall volume and number of objects increased until nine and twelve months, respectively (Table 1). This suggests a lack of expansion of the existing clusters as well as development of new clusters (Figure 2B). The volume/object ratio for Anammox and EUB on the other hand started to increase significantly after six months (t-test, p<0.005) indicating expanding volume and likely coalescence of objects (Figure 2B). Our results for old biofilm also show that within the additional year (or more) of further biofilm development the EUB and AMX cluster nearly doubled in size and number of colonies, while NOB clusters remain very stable. All three groups showed a significant correlation between the mean total volume and the mean number of objects (linear regression; R^2^_EUB_ = 0.960, R^2^_NOB_ = 0.946, R^2^_AMX_ = 0.783).

**Table 1.** Mean volume (per microscopic scan (512×512 pixel)) and mean number of objects (per microscopic scan (512×512 pixel)) derived from the FISH-CLSM analysis with DAIME. Standard deviation is based on the variation of 30 randomly analyzed pictures for every timepoint.

| | mean total volume [$\text{mm}^3$] | | | mean number of objects | | |
| --- | --- | --- | --- | --- | --- | --- |
|  | EUB | AMX | NOB | EUB | AMX | NOB |
| **1** | 6.6±4.8 | 0.10±0.3 | 0.4±0.3 | 47.4±17.2 | 2.9±7.8 | 12.1±6 |
| **3** | 11.1±7.2 | 1.5±1.5 | 1.0±1 | 83.3±21 | 30.5±13.6 | 30±19.1 |
| **6** | 25.7±17.6 | 10.2±6.8 | 4.8±5.6 | 94.9±27.5 | 66.5±29 | 63.9±31.5 |
| **9** | 44.1±35.0 | 20.5±12.6 | 6.2±3.2 | 117.6±42.5 | 63.4±24.4 | 100.6±36.6 |
| **12** | 46.4±27.2 | 36.5±23.7 | 8.2±5.4 | 138.4±37.1 | 67.8±26 | 97.8±28.7 |
| **24+** | 80.1±56.9 | 53±34.5 | 8.7±4.5 | 208.4±95 | 107.6±63.2 | 125.6±40.7 |

**Figure 2.**
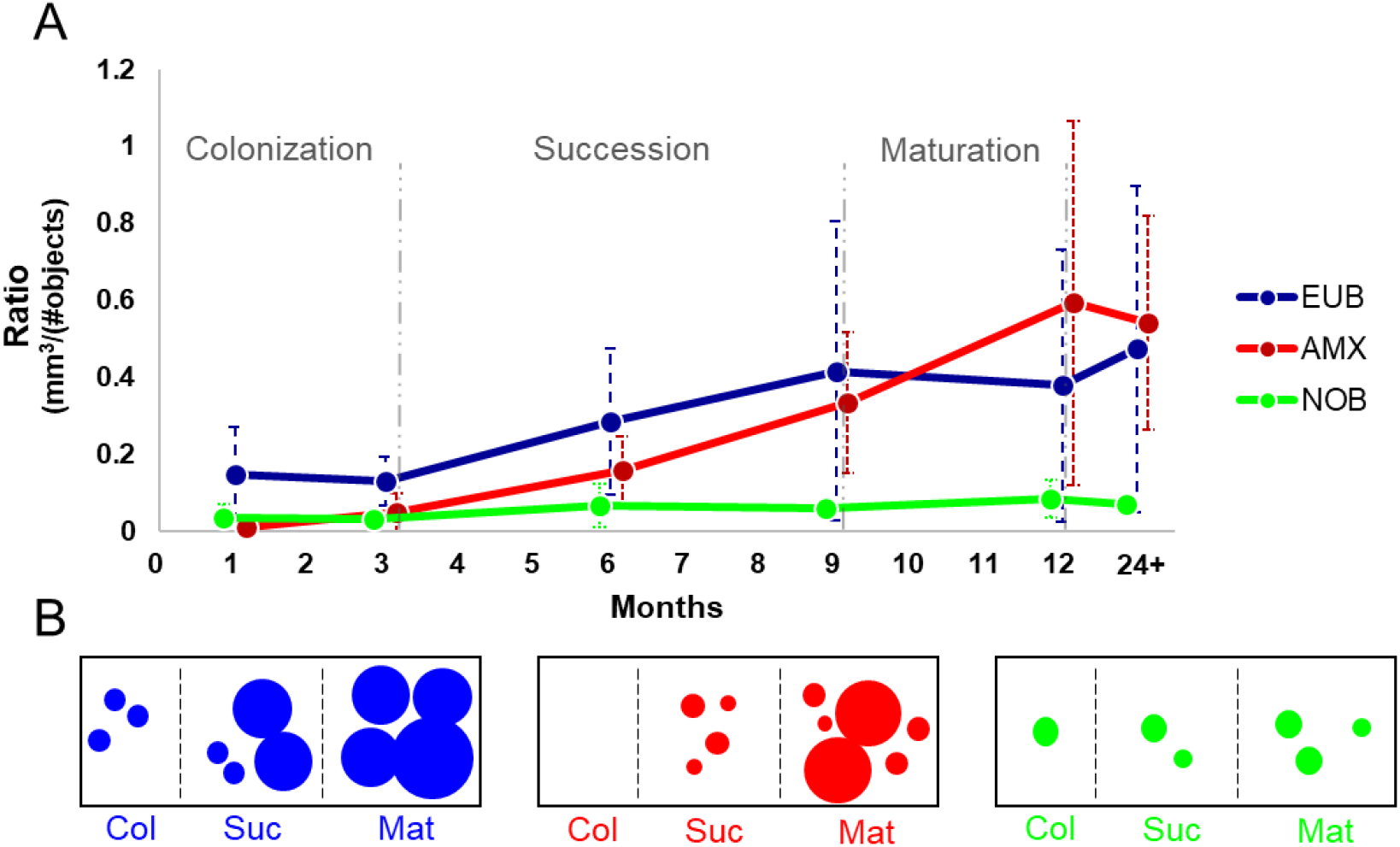
**A** The ratio between average volume and the average number of detected objects (y-axis), based on calculations from representative FISH-CLSM Image analysis, plotted over time (x-axis Colours denote stained bacterial fraction in the community: EUB = blue, NOB = green, AMX = red. Dashed lines denote the standard deviations. Dashed lines (grey) mark the distinct growth phases during biofilm formation. **B** Conceptual figure based on FISH-CLSM observations. Dot size corresponds to the volume of formed clusters, and number of detected objects.

Based on the above observations we categorized biofilm growth into three distinct biofilm growth phases. The *Colonization* (Col) phase lasts until three months, when Anammox appear and bacterial cluster volume/object ratio remains low. The *Succession* (Suc) phase includes the dynamic expansion of the bacterial and anammox clusters until month nine. The final *Maturation* (Mat) phase includes the remainder of the study period. The volume/object ratio remains mostly constant, although total biofilm volume still increases.

### Temporal dynamics of the biofilm microbial community composition

The development of the biodiversity and composition of the bacterial community were investigated by 16S rRNA gene sequencing. In total, 1038 amplicon sequence variants were detected in 102 biofilm samples. The microbial community on the biofilm carriers consisted of a diverse assemblage of taxa, many of which are typical for mainstream anammox WWTP (Bhattacharjee et al. 2017; Lawson et al. 2017; Speth et al. 2016). Throughout the observation period, the biofilm was dominated by ASVs belonging to the phyla *Proteobacteria, Bacteroidetes, Chloroflexi, Actinobacteria, Acidobacteria, Firmicutes, Planctomycetes* (primarily AMX) and *Nitrospirae* (Figure 3A). However, the distinct phases were characterized by dynamic changes in the phylum abundance structure and turnover of representatives of these phyla. During the Col phase of biofilm development ASVs that were assigned to the phyla of *Proteobacteria, Bacteroidetes, Actinobacteria* and *Firmicutes* displayed highest relative abundances. *Firmicutes* nearly vanished towards the end of the Col phase and could not reestablish their prior domination in abundance in later stages. In contrast, ASVs from the phylum *Chloroflexi* increased significantly (Students t-test, p<0.005) in their abundance, reaching up to 20% relative abundance in the Mat phase and even higher in the Old samples (Figure 3A).

**Figure 3.**
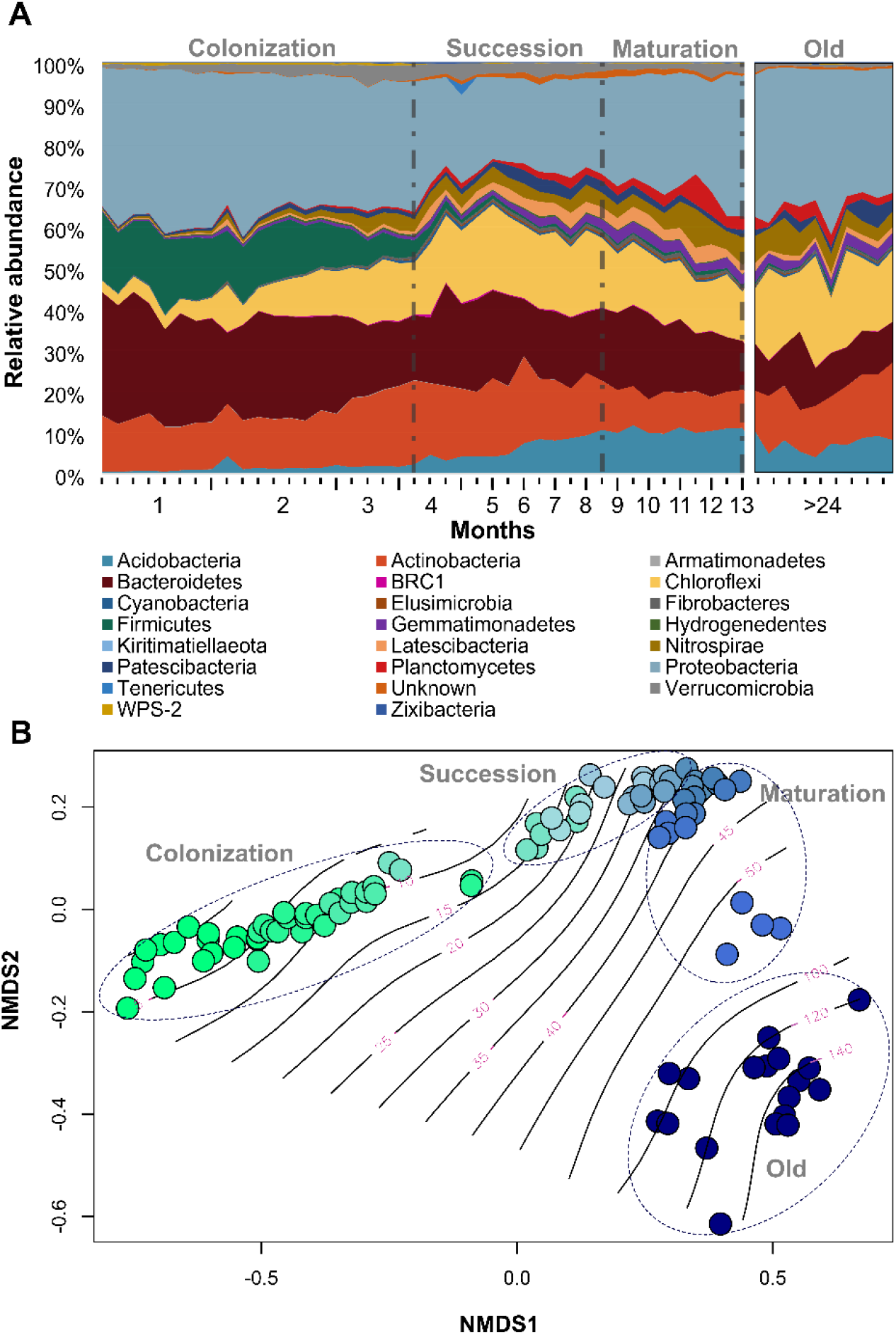
**Upper Panel:** Area chart representing the average abundance of top 15 bacterial Phyla over the year of biofilm development (n=13months) and from the old biofilms take as a reference. **Lower panel:** Non-metric multidimensional scaling analysis based on the Bray-Curtis dissimilarity between the different sampling time-points. Here, all duplicates were plotted over time. Color denotes the time: Green -> Blue. Ellipses highlight the clustering of the communities into the distinct growth phases. A GAM model (black wave lines) underscores the temporal development (5-55 Weeks and afterwards (old)). (Stress: 0.059)

Members of the *Proteobacteria* also decreased in numbers during the colonization phase, but remained the most abundant phylum in all later phases (~40%). *Actinobacteria* and *Bacteroidetes* increased during the colonization phase but decreased during succession to more or less stable abundances. This was also observed for members of the phyla *Acidobacteria* (Figure 3A). Anammox ASVs (*Planctomycetes*) and *Nitrospirae* displayed very low abundances during the first three months of biofilm growth but started to increase in numbers during the succession phase up to a maximum relative abundance of ~8% of the community. After the succession phase (nine months) the community composition stabilized with less evident fluctuations in community composition and ASV abundances (Figure 3A).

To evaluate how community structure changes over time and if our categorization into different growth phases is also reflected in the community dissimilarity, we performed a nMDS analysis on the ASV abundance data (Figure 3B). We found that dissimilarities in community composition between the three development phases as well as to the old carriers were all significantly different (Permanova, p<0.001, R^2^: 0.5783). Furthermore, community dissimilarities within categories were much more pronounced during the Col phase and gradually decreased as biofilms matured. Especially late stages of Suc and early stages of Mat (Week 30-35) did not display any significant differences (Permanova, p>0.5). Overall, community composition on the added carriers appeared to converge towards the old reference samples but remained significantly different from these over the whole course of the sampling campaign (Permanova, p<0.001). Within the old biofilms we also found significant differences (Permanova, p<0.001) between the respective sampling time points, but no clear temporal trend.

### Successional patterns of individual microbial taxa

A distribution analysis of ASVs between the distinct growth phases revealed a large number of ASVs that were abundant only during colonization. On the other hand, a larger number of ASVs were only shared between the succession and the maturation phase while being rare during the colonization phase (Supplementary Figure 1). Therefore, we classified ASVs into residents, transients and exclusives depending on their occurrence (or detection) frequencies over the previously defined growth phases and the old carriers (see Methods). We found 373 ASVs that occurred at least in 80% of samples over all phases of biofilm development and were thus classified as residents. Col and the Old phase displayed the highest number of exclusive ASVs (n=137 and 52, respectively), Suc and Mat shared the highest number of transient ASVs (n=211, late transients). Col and Suc shared 41 transient ASVs (early transients), while Col and Mat only shared 17 (late) transients.

To assess the contributions of the ASV categories to the temporal community turnover, we computed the fraction of the Bray–Curtis dissimilarity explained by the various populations (Figure 4). Overall, ASVs belonging to the resident fraction contributed most to community dissimilarity throughout (64 ± 5 %) indicating that they were highly sdynamic even though they remained in the biofilm throughout. The contribution of exsclusives to community change was highest during Col (14.7 ± 4.5 %), and interestingly again elevated in old biofilms (11.6 ± 2.9 %) compared to Suc and Mat phases (4.1 ± 1.3 and 4.6 ± 3.6%, respectively).

**Figure 4.**
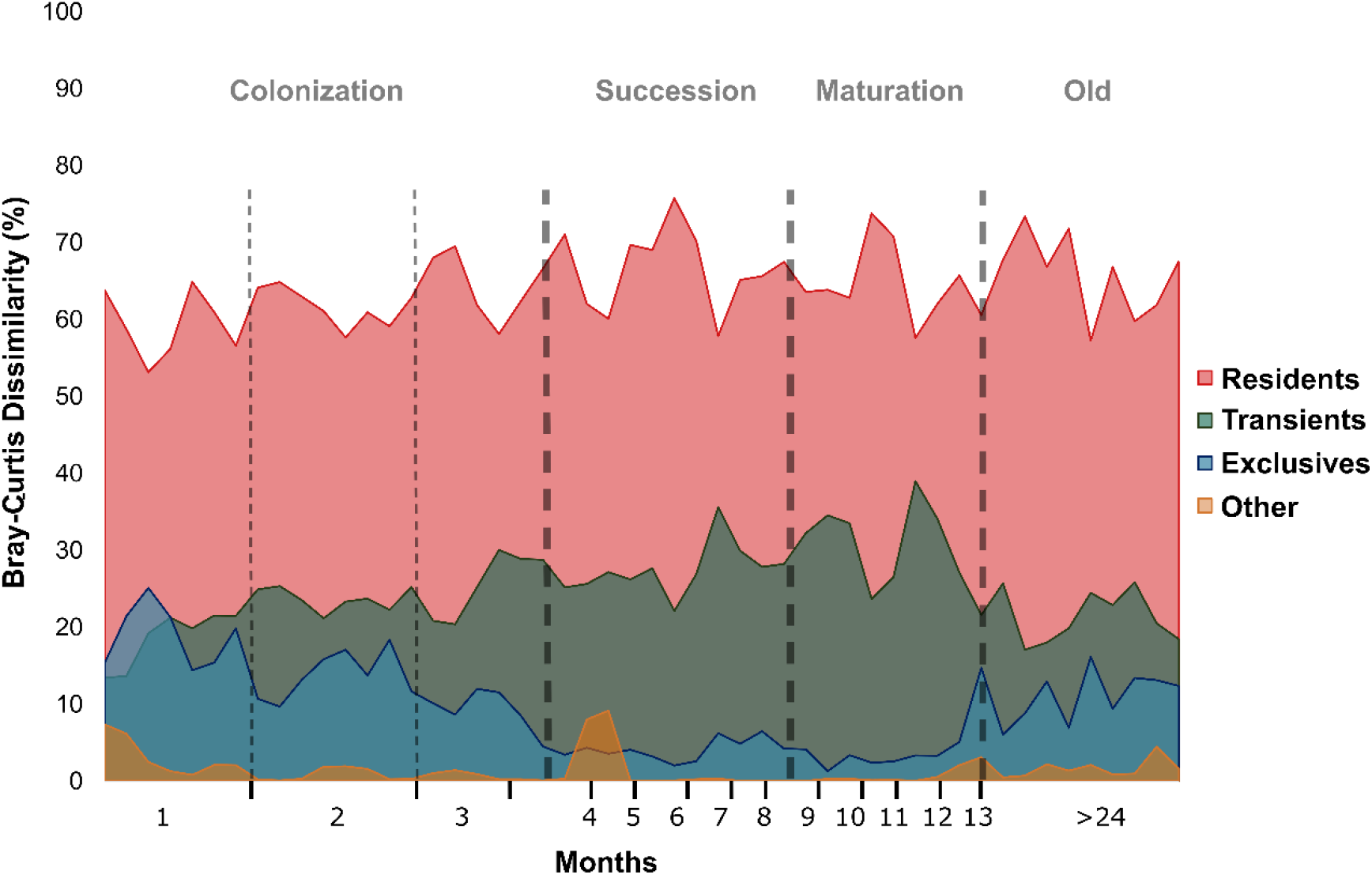
Area chart displaying the total Bray Curtis dissimilarity (100%) and average contribution of ASVs classified as Residents (red), Transients (green), Exclusives (blue) and Other (orange) during biofilm growth Month 1-13 and from the Old samples. Dashed line visually support the separation into the different growth stages. All categories sum up together to 100%.

The differences between the average contributions of exclusives were significant between all phases (Students t-test, p<0.005). The contribution of transient ASVs increased from 22 ± 3.3 % during colonization to 28 ± 3.1 % in succession and 29.7 ± 5% in the Mat phase, respectively. During the old phase, the contribution of transients decreased to 20.9 ± 3%. (Figure 4).

Pronounced community dissimilarity, high number and contribution to dissimilarity of exclusive ASVs indicated pronounced dynamics during the colonization phase. To support our findings we also calculated the average CV in abundance for all ASVs within the distinct biofilm growth phases. A high CV equals dynamic changes in the abundance of ASVs. The average CV was highest for the colonization phase (CV=1.12), while it was comparable between the succession and maturation phases (CV=0.72 & 0.82, respectively). The old biofilms were characterized by an average CV of 0.9.

### Community assembly stochasticity versus determinism during biofilm development

Next, we applied a neutral modeling approach to determine the importance of stochasticity in the community assembly. Based on the differences in the neutral model predictions (fitness of the model (r^2^), immigration rate (m)) and the NST, the colonization phase was further divided into three distinct sections (Col1, Col2, and Col3). We observed a general decrease of r^2^ and NST (from 0.75 to 0.59) over the first three months of Col (Figure 5A, B, C; Table 2) indicating initially high followed by decreasing importance of stochastic (neutral) processes.

**Table 2.** Neutral modeling of relative contribution of stochastic processes to the anammox biofilm community assembly over time. r^2^: fit of neutral model; m: immigration rate; ST: stochasticity ratio, NST: normalized stochasticity ratio

| Group | Month | neutral modeling |  | Stochasticity Ratio |  |
| --- | --- | --- | --- | --- | --- |
| | | $r^2$ | m | ST | NST |
| Col1 | 1 | **0.9108** | 0.8618 | 0.8477 | **0.7500** |
| Col2 | 2 | **0.8878** | 0.4912 | 0.8529 | **0.7378** |
| Col3 | 3-4 | **0.7434** | 0.4074 | 0.7691 | **0.5940** |
| Suc | 6-8 | **0.9044** | 1.1074 | 0.8754 | **0.7794** |
| Mat | 10-13 | **0.7264** | 0.6086 | 0.8880 | **0.8171** |
| Old | 24+ | **0.8293** | 0.8531 | 0.8439 | **0.7425** |

**Figure 5.**
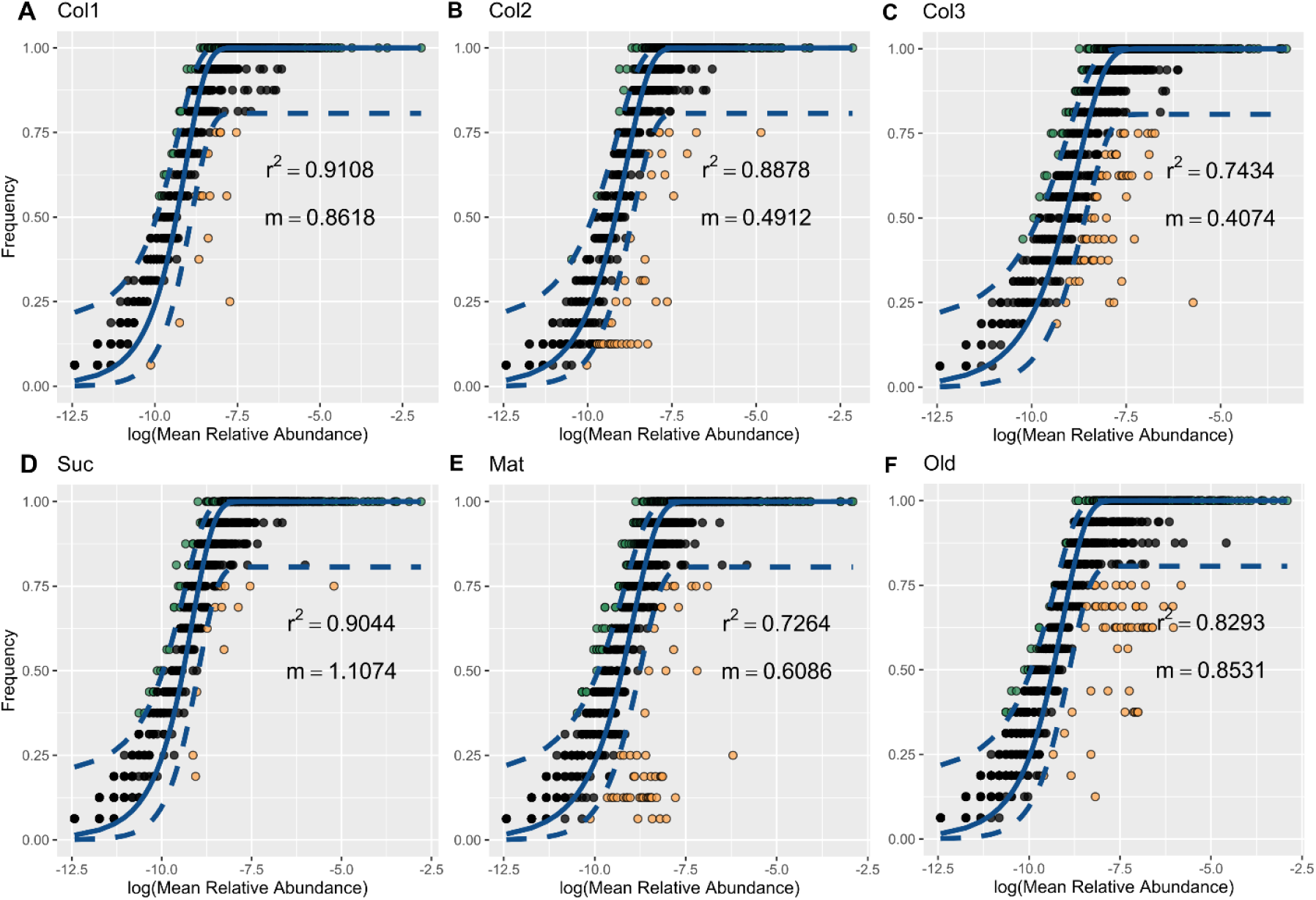
Neutral biofilm assemblage model. The analysis was conducted using normalized abundance matrix of ASVs (i.e., 100% ASVs, proxy for species) in 48 samples (16 sample each stage) collected over time. A-F, Neutral modeling plots showing the fit of neutral model, r^2^, and the migration rate, m; neutral model predictions and its corresponding 95% confidence intervals (lines) and actual distributions (points) across the stage. Colors denote prediction level: Blue:fit; Green:above; Yellow:below

Immigration rate also decreased indicating a decreasing contribution of the arrival of new taxa. The succession phase on the other hand went along with an increase of the corresponding r^2^, NST (0.9 and 0.77, respectively) and m (1.1) in comparison to Col3 (Figure 5D; Table2) indicating a resurgence of neutral processes and invasion of the developing biofilm by new taxa. In the Mat phase (Figure 5E; Table 2) r^2^ and m drastically decreased once again (0.73 and 0.60 respectively), while the stochasticity ratio increased further to 0.81. The Old carriers (Figure 5F; Table2) displayed an average fit of r^2^ = 0.83 and the second highest immigration rate (0.85). For the old carriers we calculated an average NST of 0.74.

### Dramatically changing deterministic co-occurrence and species segregastion patterns

We further explored co-occurrence patterns using network inference from strong and significant correlations of the Spearman’s rank coefficient. Resulting networks for each of the anammox growth stages differed substantially in number of nodes (ASVs), edges (number of connections), size of modules (closely interconnected nodes), modularity (extent to which species interaction are organized into modules or subnetworks in a network) and the clustering coefficient (the degree to which nodes tend to cluster together) (Figure 6; Table 3). In all parameters except modularity, we observed a decrease from Col1 to Col2, whereas all network properties significantly increased again from Col2 to Col3 (Figure 6A, B, C; Table 3). The succession phase displayed lowest values for all network properties with an all-time low in nodes (n=94) and edges (n=87) but an increase in average modularity from 0.295 (Col3) to 0.795 (Figure 6D). During maturation the number of nodes and edges increased again (n=213 and 206 respectively). In addition, an increase in modularity was observed (Figure 6F; Table 3). Networks for Old carriers that have been in the reactor for over two years displayed again a substantial increase in nodes (n=363) and edges (N=1027) but decrease in modularity (0.687) in comparison to the Mat biofilm.

**Table 3.**
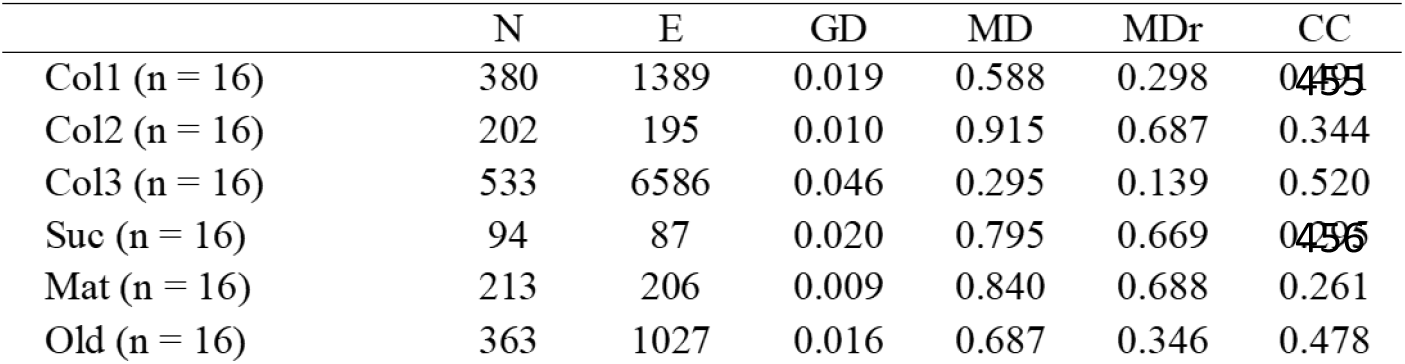
Comparison of topological properties of observed co-occurrence networks of AMX biofilm communities with identically sized Erdos-Reyni random networks. Abbreviations: N, number of nodes; E, number of edges; GD: graph density; MD: modularity, CC: cluster coefficient. Subscript r indicates the property of the random networks

**Figure 6.**
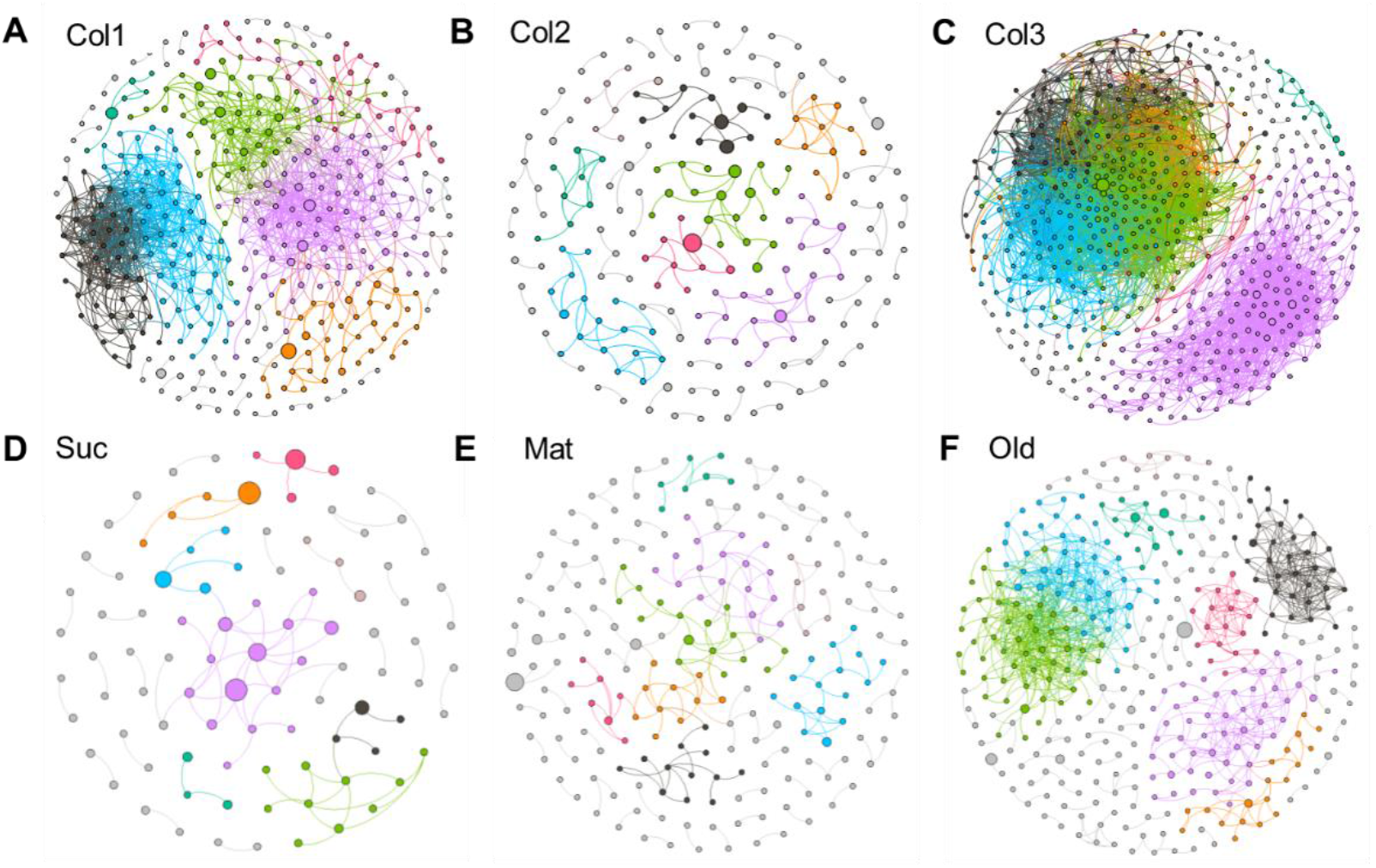
Co-occurrence model of AMX biofilm community assembly. A-F Co-occurrence networks colored by modularity class. Each node represents a species with its size proportional to its relative abundance in all samples, and each edge stands for a very strong (Spearman’s ρ ≥ 0.8) and significant (FDR-adjusted P-value < 0.05) Spearman’s correlation. More information see Table 3 and Material and Methods.

Analyzing the networks for co-occurrence of ASV within and between individual Phyla, we found that in Col1 ASVs tended to not co-occur deterministically as shown by low O/R ratios (Supplementary Table 2). Only ASVs assigned to the phylum of *Bacteroidetes* tended to significantly co-occur with other representatives of the same phylum (intra-taxon co-occurrence) during this phase. This changed drastically in Col2 and Suc, where the number of significant positive inter-taxa and intra-taxon co-occurrences increased substantially. Especially the Suc phase of biofilm formation displayed a pronounced number of inter-taxa co-occurrences. In stark comparison, during the Col3, Mat and Old phases incidences of inter- and intra-taxa co-occurrences tended to occur less than would be expected by chance.

Based on the findings from the co-occurrence networks, we applied a null-model approach to make inferences about the ecological species interactions (cooperation and competition) within the microbial community over the growth phases of the biofilm. Here, we found strong consistencies between the C-score and its variance (Cvar-score) (Table 4, Supplementary Figure 2). During colonization SES values from C-score increased from 3.32 to 45.3 while the corresponding Cvar-score increased from 3.11 to 285, suggesting increasingly less positive species interaction (cooperation) over time while species competition increased (Table 4, Supplementary Figure 2). SES values and potential species competition decreased for both scores during the succession phase followed by a significant increase in the mature phase (SES: 34.6 and 46.8, respectively). Species cooperation increased again in old carriers in comparison to the mature phase with comparable SES values from 17.8 (C-score) and 14.8 (Cvar-score), respectively (Table 4, Supplementary Figure 2).

**Table 4.**
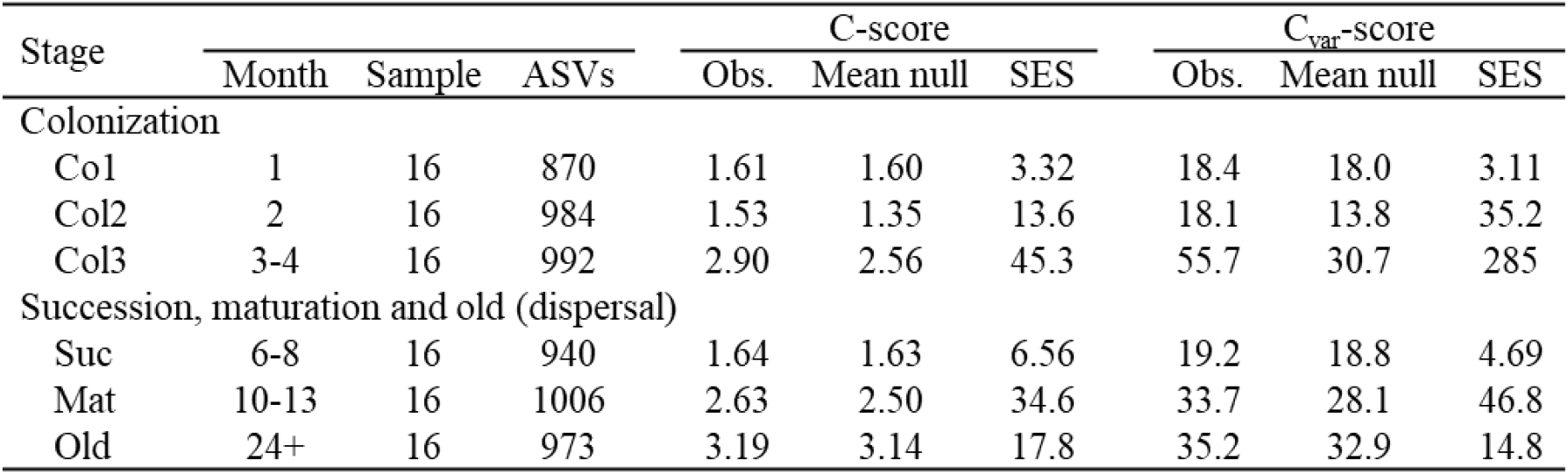
Observed C-scores and C_var_-scores, mean metric values under null models, standardized effect sizes (SES) for annamox biofilm communities over time. Higher observed SES values of C-score suggest greater degrees of species segregation than would be expected by chance. Higher SES values of C_var_-score indicate greater degrees of both species segregation and aggregation. All *P*-values are < 0.0001.

| Stage | | | | C-score | | | $C_{\text{var}}\text{-score}$ | | |
| --- | --- | --- | --- | --- | --- | --- | --- | --- | --- |
|  | Month | Sample | ASVs | Obs. | Mean null | SES | Obs. | Mean null | SES |
| Colonization |  |  |  |  |  |  |  |  |  |
| Col1 | 1 | 16 | 870 | 1.61 | 1.60 | 3.32 | 18.4 | 18.0 | 3.11 |
| Col2 | 2 | 16 | 984 | 1.53 | 1.35 | 13.6 | 18.1 | 13.8 | 35.2 |
| Col3 | 3-4 | 16 | 992 | 2.90 | 2.56 | 45.3 | 55.7 | 30.7 | 285 |
| Succession, maturation and old (dispersal) |  |  |  |  |  |  |  |  |  |
| Suc | 6-8 | 16 | 940 | 1.64 | 1.63 | 6.56 | 19.2 | 18.8 | 4.69 |
| Mat | 10-13 | 16 | 1006 | 2.63 | 2.50 | 34.6 | 33.7 | 28.1 | 46.8 |
| Old | 24+ | 16 | 973 | 3.19 | 3.14 | 17.8 | 35.2 | 32.9 | 14.8 |

## Discussion

The spatial and successional dynamics during biofilm formation of microbial communities on suspended sludge or granules within AMX reactor systems were the focus of recent research (Wang et al. 2019; Chu et al. 2015). However, given the importance and advantages of carrier attached biofilms in engineered systems and the distinct microbial community succession on biofilms (Niederdorfer, Peter, and Battin 2016) it is crucial to investigate the ecological processes that lead to functional established AMX biofilms in waste water systems. This study was designed to track and model AMX biofilm microbiota development on synthetic carrier material in a controlled engineered ecosystem over the course of a year. Importantly, the presence of a surfeit of mature anammox biofilm carriers in the reactor is thought to have largely eliminated the importance of dispersal limitation and random founder effects. This allows a clear view on the internal dynamics of the colonization process.

In accordance with our hypothesis ii) and in agreement with the fundamentals of biofilm research we found that distinct developmental stages in biofilm formation emerged naturally from the data in terms of structure (Figure 1, 2) and community composition (Figure 3A). In support of hypothesis iii) these stages were further characterized by dynamic shifts between stochastic and deterministic processes (Figure 5) and changing co-occurrence patterns (Figure 6), until converging into community structures that resembled the mature carrier biofilms already present in the reactor (Figure 3B). In the following sections we discuss the ecological dynamics at each stage with Figure 7 as a conceptual guideline.

**Figure 7.**
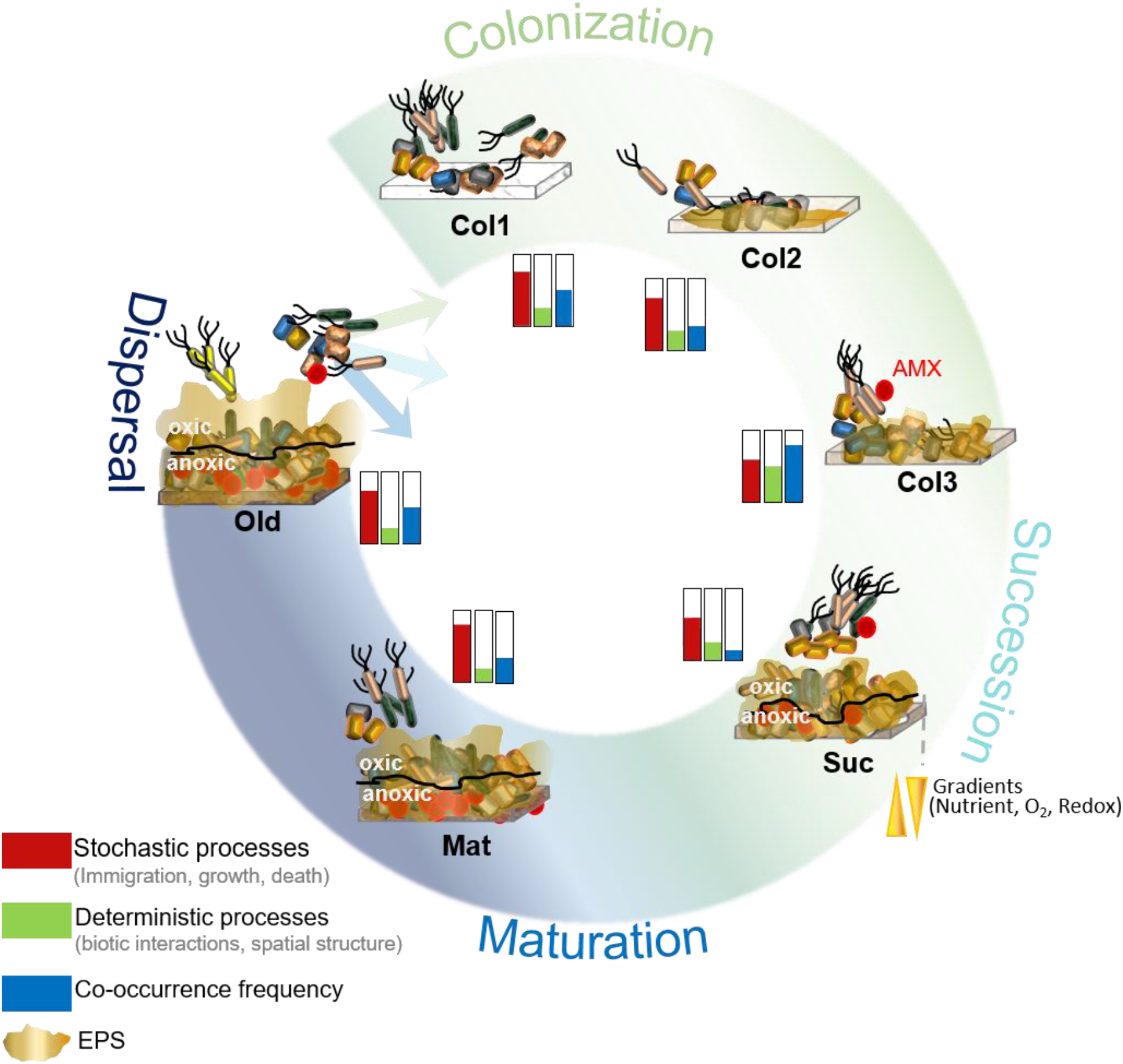
Conceptual framework of the biofilm microbiota development on synthetic carrier material. Starting clockwise from Colonization to Maturation with the continuous dispersal from the old Carriers at the end. Barcharts denote relative contributions of ecological parameters during each observed development stages. (Red: stochastic processes, Green: Deterministic processes and Blue: Co-occurrence frequencies). Amount of bacteria swimming towards the biofilm is relative to the modelled immigration rates. The black line denotes the oxic/anoxic interface within the biofilm.

### Initial colonization phase (Col1)

In support of our hypothesis i) biofilm formation was initiated by a consortium of heterotrophic bacteria with mostly members of *Firmicutes*, *Bacteroidetes*, *Proteobacteria* and *Actinobacteria* (Figure 3A). Using FISH-CLSM (Figure 1), we observed a stochastic spatial distribution of microcolonies built by pioneering colonizers in the first month. Presumably, detached bacterial cells from mature biofilms or from the surrounding wastewater non-specifically and randomly attached to new carrier surfaces. As newly adherent cells are loosely associated with carriers, they are likely to detach again as they are subjected to hydrodynamic perturbation (e.g., shear forces) in the reactor (Figure 1). This is reflected by the substantially higher quantity of exclusive ASVs at the very beginning of biofilm formation (Figure 4) and a higher variation in abundance (CV). Furthermore, neutral modeling revealed the predominant role of stochastic processes in the microbiota assembly during this initial colonization stage (Col1) (Figure 5A, Table 2). However, a large fraction of the initial founding taxa established successfully and became part of the biofilm core community. All observed initial colonization patterns during Col1 generally agree with the scientific consensus on first steps of biofilm formation (Hall-Stoodley, Costerton, and Stoodley 2004; Jackson, Churchill, and Roden 2001; Battin et al. 2016).

### Later colonization phases (Col2 & Col3)

During Col2, stochastic processes still dominated the biofilm formation, but the relative contribution of deterministic processes increased due to higher competition (Figure 5B, Table 4). We argue that the microscopically observed stagnation in cluster formation (Figure 1, Month 3) might directly result from increased competition, resulting in a metabolic trade-off (Polz and Cordero 2016), where pioneering populations have to invest their metabolic capacity rather into defense mechanisms and the production of extrapolymeric substances (EPS) than in fast growth and spatial expansion. The increase in determinism (competition) accompanied by a considerable decrease of immigration rate further supports the notion of the community transitioning from initial reversible to irreversible attachment with the result of enhanced community stability. The drastic decrease in exclusive ASVs in Col3 (Figure 4) supports this result. While pioneering Phyla like *Firmicutes* slowly vanished, new key players, such as *Chloroflexi* emerged (Figure 3A). We argue that a credible mechanism that may underlie these dynamics is substrate utilization. During initial biofilm colonization, labile substrates will be consumed first, supporting copiotrophic microbial taxa, such as Firmicutes (Datta et al. 2016; Nemergut et al. 2016). These are later replaced by more oligotrophic (slow growing) taxa (e.g. *Chloroflexi*, *Acidobacteria)* that are capable of metabolizing the remaining, more recalcitrant, organic C pools (Rui, Peng, and Lu 2009) and potentially outcompete less adapted pioneering consortia (Polz and Cordero 2016). The filamentous members of the Chloroflexi phylum are known to structurally support biofilm formation (Speirs et al. 2019) and to scavenge secondary metabolites (Kindaichi et al. 2012; Xia, Kong, and Nielsen 2007). These ongoing dynamics in community turnover are also reflected in the dissimilarity analysis (Figure 3B) and agree with the significantly increased species competition (Fig. 5C) and frequencies of species co-occurrence associations (Fig. 6C), revealing a substantial increase in the deterministic influences on the microbiota assembly.

### Succession phase (Suc)

Upon the accomplishment of irreversible attachment, cells multiply and continue producing essential EPS matrix components, forming microcolonies and eventually multilayer aggregates (Hall-Stoodley, Costerton, and Stoodley 2004). The expansion of the clusters in the carrier material (Figure 1, Table 1 (Month 6-9)) are supporting this notion. Additionally, the increasing number of late transient ASVs (Figure 4, Supplementary Figure 1) and the reduced community dissimilarities (Figure 3B) point into the direction of convergence towards a stable core community. Understood in the sense that the competition-driven, deterministic dynamics have now played out, this interpretation is in line with the Suc phase also being characterized by a sharp increase in both the importance of stochastic assembly and immigration rate (Figure 5D, Table 2) and by decreased strengths of species competition and co-occurrence (Figure 6D and Supplementary Figure 3). A stable core community now enables transitory attachment and immigration events to dominate the temporal dynamics again.

During the Suc phase, AMX first started to emerge within the community (Figure 1, 3A) which confirmed our hypothesis i) that AMX do not participate in the initial biofilm formation but rather grow slowly into the biofilm when conditions are sufficiently beneficial. Actively respiring aerobic microcolonies can consume oxygen faster than it diffuses through the biofilm, which results in the formation of anaerobic zones in lower layers of the biofilm, whereas upper layers remain aerobic (Flemming et al. 2016). We argue, that slow growth rates (Mulder et al. 1995; Kuenen 2008) and sensitivity to various environmental factors (Jin et al. 2012) make anammox bacteria weak competitors during initiation of biofilm formation. However, the formation of specific niches at the oxic/anoxic interface and their autotrophic lifestyle enables them to increase in biomass once a biofilm has established (Zhang and Okabe 2020). During biofilm development complex physical and chemical gradients (available nutrients, oxygen, redox gradients) are established (Veach et al. 2016) and create multiple niches that harbor various cells with different metabolic capacities, which in turn creates opportunities for cooperation (Flemming et al. 2016). We observed an increase in inter-taxa co-occurrences between taxonomically distant bacteria (Supplementary Table 2) suggesting a high degree of mutualistic and commensal metabolic interactions are shaping the community during the Suc phase (Ju and Zhang 2015).

### Mature phase (Mat & Old)

The near complete filling of the pore space with biomass, and the agglomeration of existing and formation of new clusters (Figure 1, 2) along with the stability in community structure (Figure 3, 4) reflect the definition of a stable and mature biofilm. Bacterial cells synthesize and release signaling molecules (e.g., c-di-GMP) to sense the presence of each other, promoting e.g. production of EPS matrix (Romling, Galperin, and Gomelsky 2013; Rinaldo et al. 2018; McDougald et al. 2012) while expanding and maintaining the mature biofilm. We speculate that the observed increase in modularity (Table 3) could be a direct result of the released signal molecules that led to the creation of new ecological niches and agglomeration of the clusters (Figure 1). The expansion of the biofilm along with increasing EPS production greatly increases the potential interactions between biofilm inhabitants. Given the high complexity of EPS molecules, their degradation requires multiple enzymes (Flemming and Wingender 2010), which in turn creates multiple opportunities to foster species interactions through cross-feeding or co-degradation (Nadell, Drescher, and Foster 2016). For AMX consortia in reactor ecosystems it has already been shown that cross-feeding between multiple members of the microbial community potentially improves the anammox metabolism (Lawson et al. 2017; Yunpeng Zhao et al. 2018).

During Mat phase and in Old biofilm intra-taxa co-occurrences further increased, which suggests established niches and an increase in the co-occurrence of closely related species (Röttjers and Faust 2018). According to the established concept of the biofilm life cycle, bacterial cells disperse from mature biofilm, revert to planktonic growth and start a new cycle of biofilm establishment on a new substrate (McDougald et al. 2012; Flemming et al. 2016). In engineered ecosystems, the passive sloughing caused by shear forces introduced by constant stirring is the predominant reason of biofilm dispersal (Garny, Horn, and Neu 2008) and increases with increasing biofilm thickness (Telgmann, Horn, and Morgenroth 2004). We argue that these dispersal dynamics also create new ecological niches within the biofilm community, which are susceptible to attachment by scavenging bacterial cells (McDougald et al. 2012). The observed increase in immigration rate, number of exclusive ASVs and frequency of co-occurrences (Figure 6F) support this notion. On the other hand, the marked decrease in the community-wide species competition and increased influence of stochastic processes (Figure 5F, Table 2, Supplementary Figure 2) on the community assembly might be a direct result of the aforementioned dispersal dynamics.

In summary, our findings show in considerable detail how an AMX biofilm develops on synthetic carrier material within an engineered ecosystem. Stochastic and deterministic processes structure the microbial community, and lead to a multi-phase succession even in the presence of abundant established biofilms in the reactor. The right choice of reactor conditions has proven its potential to facilitate AMX growth within biofilms (Agrawal et al. 2017; Tao et al. 2012; Bhattacharjee et al. 2017). Our results suggest, however, that the microbial community has to proceed through the different phases of biofilm formation mandated by internal ecological processes to reach the desired core community structure and metabolic capacities of the mature biofilm. Simple changes in reactor conditions alone are thus unlikely to accelerate or modulate biofilm formation as the community members themselves, their mutual interactions and the biofilm structure have to develop over time (in our case over a year) and alter the microenvironment they inhabit. This has considerable practical implications. It may prove difficult to considerably speed up the process of establishing functional AMX biofilms as each phase depends on previous developments, which puts constraints on the startup of new reactors and the regular replacement of carrier material. Further, given the observed high degree of microbial interactions in AMX reactors ((Wang et al. 2019; Lawson et al. 2017) and Fig. 6) it seems possible that changes in the reactor configuration might alter the community development, which in turn could directly affect the balance of bacterial interactions within the community. AMX bacteria generally comprise only a small fraction of the microbial community and are, especially under mainstream conditions, sometimes very susceptible to disturbances. However, they are the key players for one of the most promising energy-efficient mechanisms of fixed nitrogen elimination and the effect of changing community interactions on AMX ecology thus needs to be further explored. On the other hand, our results indicate that the developing biofilm on new carriers converged towards the community composition and structure observed similar to already established biofilms. The mature biofilm had considerable long-term stability both in terms of function and microbial composition, which may point to a stabilizing effect of the complex interactions in the diverse microbial biofilm community. A deeper understanding of the functions involved in the assembly and maintenance of AMX biofilms is clearly necessary to better understand and potentially manipulate their development.

## Acknowledgements

This work was supported by funding from the Swiss national science foundation Sinergia project ISOMOL: CRSII5_170876. The authors would like to thank Moritz Lehmann (University of Basel) and Joachim Mohn (EMPA) for the helpful scientific discussions during the course of this study. We would like to thank the ScopeM facility and employees of the ETH Hönggerberg for providing access to their infrastructure and support for cryosectioning and microscopic analysis. We would also like to thank the technical staff of the Versuchshalle EAWAG in Dübendorf and the GDC for providing access to and support of bioinformatics analysis performed on the ETH Zurich Euler cluster. Furthermore, we would like to thank the MicEcol group in Kastanienbaum for the fruitful manuscript discussions.

## Author contributions

R.N, L.F and H.B designed the study. All authors provided helpful feedback and suggestions throughout work on the study. R.N and D.H. performed the experiment. L.F and D.H performed the sampling. R.N and L.F performed the laboratory work, sequencing and data analysis. L.F performed FISH-CLSM and entailed analysis. L.Y and F.J performed all modelling approaches and co-occurrence analysis. R.N wrote the first draft of the manuscript based on the master thesis by L.F. The manuscript was written by R.N, H.B and F.J with critical and helpful reviews from D.H, J.W, L.Y, P.M, A.J and L.F.

## Additional information

Competing interests: The authors declare no competing financial interest

## Data Availability

Raw 16S sequences can be found on the NCBI sequence read archive under the repository number: PRJNA636402

All other data (Gene abundance tables as comma separated tables and R codes) are available from the Eawag Research Data Institutional Collection (Eric) at [URL to be supplied after acceptance of manuscript].

