## Supplementary material for "Distinct growth stages shaped by an interplay of deterministic and neutral processes are indispensable for functional anammox biofilms"

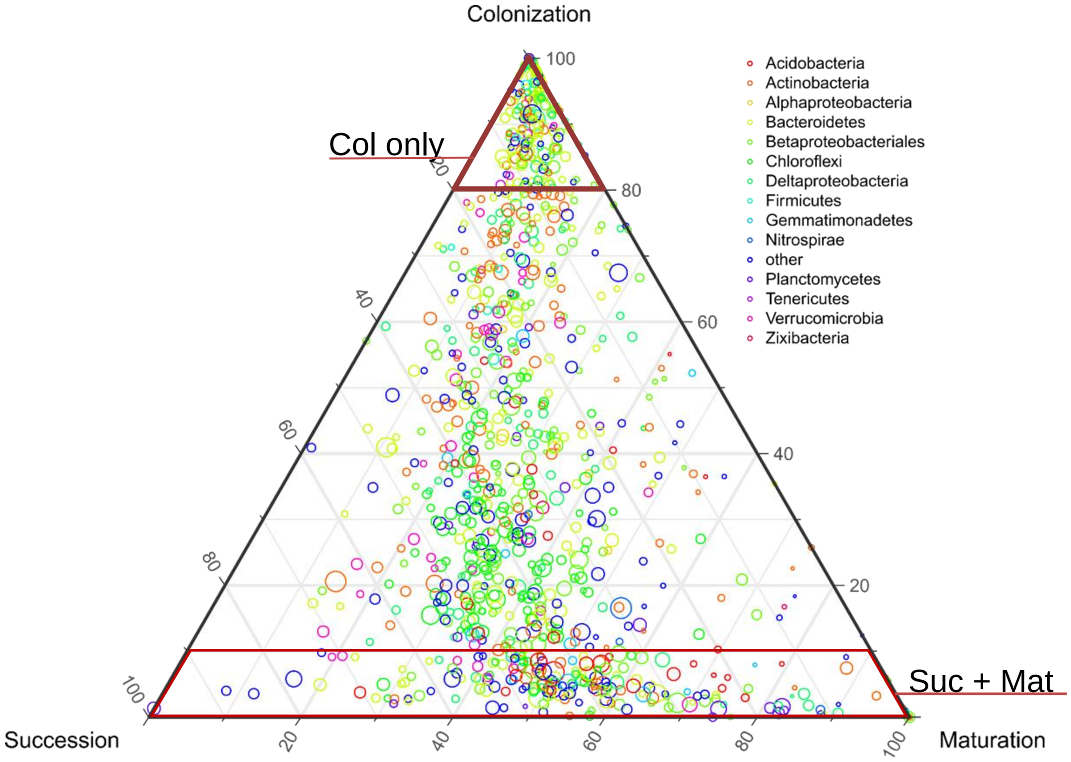


**Supplementary Figure 1** Ternary plot based the relative abundance The position in the triangle indicates the relative abundance of each taxon among the three development phases; the size of the circle represents the relative abundance of taxa. Red shapes highlight ASVs only occurring in the Col phase and the shared ASVs between Suc and Mat.

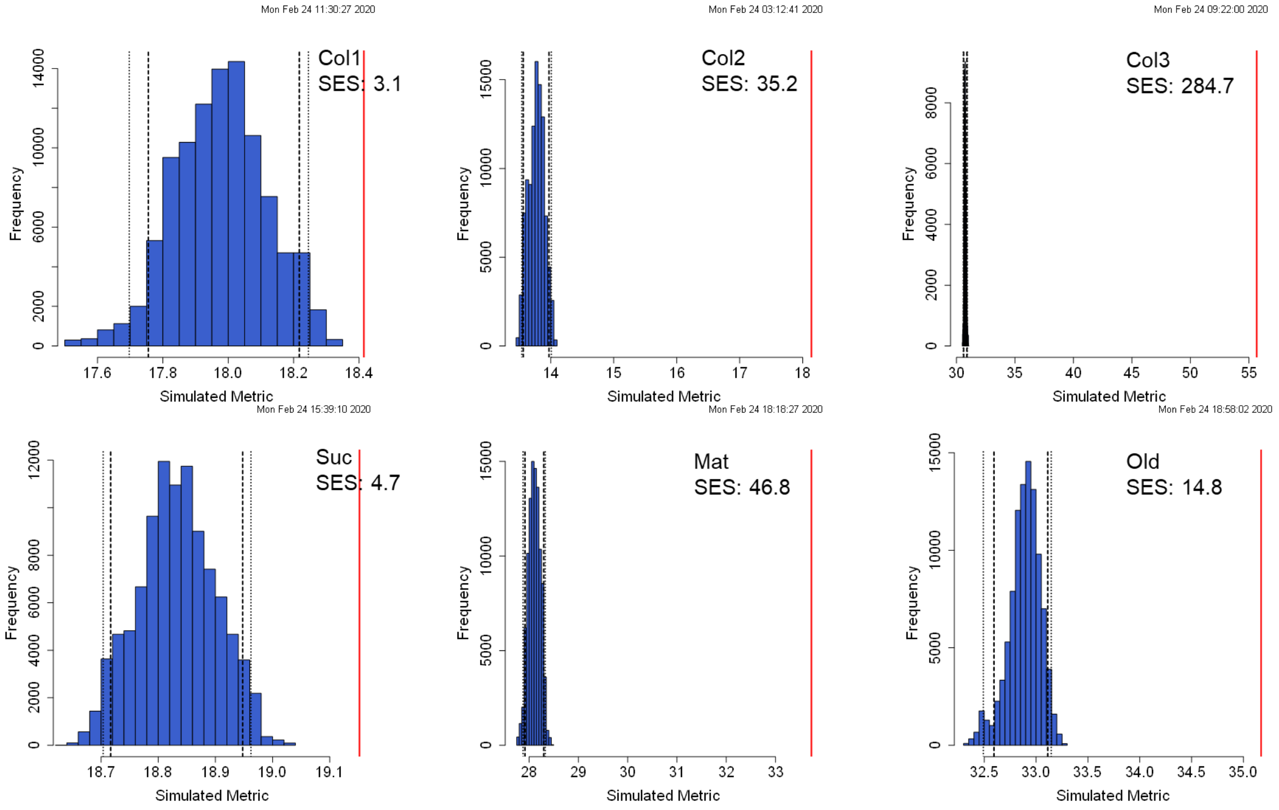


**Supplementary Figure 2** Observed C-score$_{var}$ (red lines) and the distributions of C-score$_{var}$ expected by null models (histograms) which assume no species interactions. A red line to the right of a histogram denotes fewer co-occurrences between species in that community than would be expected by chance (P-values <0.0001). The dash lines indicate 1-tail and 2-tail 95% confidence intervals of the mean C-score of the of the null models. Higher SES values of C$_{var}$-score indicate greater degrees of both species segregation and aggregation.

**Table S1|** List of oligonucleotide probes used in this study. FISH-probes are labeled with fluorescent dyes. Cy = cyanine, FITC = fluorescein isothiocyanate.

| Probe | Target organisms | Label | Sequence (5' → 3') | Source |
|---|---|---|---|---|
| AMX820 | Ca. *Brocadia* , Ca. *Kuenenia* | Cy3 | AAA ACC CCT CTA CTT AGT GCC C | Schmid et al., 2000 |
| EUB338 | Most bacteria | Cy5 | GCT GCC TCC CGT AGG AGT | Amann et al., 1990 |
| EUB338 II | Phylum *Planctomycetales* | Cy5 | GCA GCC ACC CGT AGG TGT | Daims et al., 1999 |
| EUB338 III | Phylum *Verrucomicrobiales* | Cy5 | GCT GCC ACC CGT AGG TGT | Daims et al., 1999 |
| NON338 | no target (control probe) | Cy5 | ACT CCT ACG GGA GGC AGC | Wallner et al., 1993 |
| Ntspa662 | Genus *Nitrospira* | FITC | GGA ATT CCG CGC TCC TCT | Daims et al., 2001 |
| Ntspa712 | Most of Phylum *Nitrospirae* | FITC | CGC CTT CGC CAC CGG CCT TCC | Daims et al., 2001 |
| NIT3 | Nitrobacter sp. | FITC | CCT GTG CTC CAT GCT CCG | Wagner et al., 1996 |

**Table S2 Phylum-level observed and random incidences of co-occurrence association** Orange color highlights the O/R ratios above 1.0 and therefore higher than expected by chance

| Stage | Node Type 1 (T1) | Node Type 2 (T2) | #T1 | #T2 | Edges | R% | O% | O/R-ratio |
|---|---|---|---|---|---|---|---|---|
| Network All: 183 nodes and 688 edges | | | | | | | | |
| | *p_Acidobacteria* | *p_Latescibacteria* | 11 | 4 | 7 | 0.26 | 1.02 | 3.85 |
| | *p_Acidobacteria* | *p_Acidobacteria* | 11 | 11 | 8 | 0.33 | 1.16 | 3.52 |
| | *p_Actinobacteria* | *p_Firmicutes* | 34 | 3 | 11 | 0.61 | 1.60 | 2.61 |
| | *p_Bacteroidetes* | *p_Planctomycetes* | 42 | 2 | 9 | 0.50 | 1.31 | 2.59 |
| | *p_Firmicutes* | *p_Proteobacteria* | 3 | 56 | 15 | 1.01 | 2.18 | 2.16 |
| | *p_Acidobacteria* | *p_Bacteroidetes* | 11 | 42 | 39 | 2.77 | 5.67 | 2.04 |
| | *p_Patescibacteria* | *p_Proteobacteria* | 5 | 56 | 5 | 1.68 | 0.73 | 0.43 |
| | *p_Actinobacteria* | *p_Patescibacteria* | 34 | 5 | 3 | 1.02 | 0.44 | 0.43 |
| | *p_Actinobacteria* | *p_Chloroflexi* | 34 | 14 | 8 | 2.86 | 1.16 | 0.41 |
| | *p_Bacteroidetes* | *p_Chloroflexi* | 42 | 14 | 9 | 3.53 | 1.31 | 0.37 |
| | *p_Chloroflexi* | *p_Proteobacteria* | 14 | 56 | 10 | 4.71 | 1.45 | 0.31 |
| Network Col1: 380 nodes and 1389 edges | | | | | | | | |
| | *p_Bacteroidetes* | *p_Bacteroidetes* | 100 | 100 | 346 | 6.87 | 24.91 | 3.62 |
| | *p_Gemmatimonadetes* | *p_Proteobacteria* | 6 | 152 | 8 | 1.27 | 0.58 | 0.45 |
| | *p_Bacteroidetes* | *p_Verrucomicrobia* | 100 | 13 | 11 | 1.81 | 0.79 | 0.44 |
| | *p_Actinobacteria* | *p_Chloroflexi* | 52 | 26 | 11 | 1.88 | 0.79 | 0.42 |
| | *p_Others* | *p_Proteobacteria* | 9 | 152 | 11 | 1.90 | 0.79 | 0.42 |
| | *p_Bacteroidetes* | *p_Chloroflexi* | 100 | 26 | 14 | 3.61 | 1.01 | 0.28 |
| | *p_Chloroflexi* | *p_Proteobacteria* | 26 | 152 | 20 | 5.49 | 1.44 | 0.26 |
| Network Col2: 202 nodes and 195 edges | | | | | | | | |
| | *p_Armatimonadetes* | *p_Verrucomicrobia* | 2 | 13 | 4 | 0.13 | 2.05 | 16.02 |
| | *p_Verrucomicrobia* | *p_Verrucomicrobia* | 13 | 13 | 10 | 0.38 | 5.13 | 13.35 |
| | *p_Planctomycetes* | *p_Verrucomicrobia* | 3 | 13 | 3 | 0.19 | 1.54 | 8.01 |
| | *p_Firmicutes* | *p_Actinobacteria* | 2 | 22 | 3 | 0.22 | 1.54 | 7.10 |
| | *p_Planctomycetes* | *p_Actinobacteria* | 3 | 22 | 2 | 0.33 | 1.03 | 3.15 |
| | *p_Chloroflexi* | *p_Chloroflexi* | 31 | 31 | 14 | 2.29 | 7.18 | 3.13 |
| | *p_Actinobacteria* | *p_Others* | 22 | 7 | 4 | 0.76 | 2.05 | 2.70 |
| | *p_Others* | *p_Verrucomicrobia* | 7 | 13 | 2 | 0.45 | 1.03 | 2.29 |
| | *p_Planctomycetes* | *p_Chloroflexi* | 3 | 31 | 2 | 0.46 | 1.03 | 2.24 |
| | *p_Verrucomicrobia* | *p_Actinobacteria* | 13 | 22 | 1 | 1.41 | 0.51 | 0.36 |
| | *p_Others* | *p_Bacteroidetes* | 7 | 45 | 1 | 1.55 | 0.51 | 0.33 |
| | *p_Chloroflexi* | *p_Proteobacteria* | 31 | 67 | 5 | 10.23 | 2.56 | 0.25 |
| | *p_Verrucomicrobia* | *p_Bacteroidetes* | 13 | 45 | 1 | 2.88 | 0.51 | 0.18 |
| Network Col3: 533 nodes and 6586 edges | | | | | | | | |
| | *p_Actinobacteria* | *p_Patescibacteria* | 77 | 10 | 87 | 0.54 | 1.32 | 2.43 |
| | *p_Actinobacteria* | *p_Verrucomicrobia* | 77 | 24 | 35 | 1.30 | 0.53 | 0.41 |
| | *p_Actinobacteria* | *p_Chloroflexi* | 77 | 52 | 70 | 2.82 | 1.06 | 0.38 |
| | *p_Planctomycetes* | *p_Proteobacteria* | 9 | 186 | 28 | 1.18 | 0.43 | 0.36 |
| | *p_Proteobacteria* | *p_Verrucomicrobia* | 186 | 24 | 66 | 3.15 | 1.00 | 0.32 |
| | *p_Bacteroidetes* | *p_Verrucomicrobia* | 118 | 24 | 41 | 2.00 | 0.62 | 0.31 |
| | *p_Bacteroidetes* | *p_Chloroflexi* | 118 | 52 | 64 | 4.33 | 0.97 | 0.22 |
| | *p_Chloroflexi* | *p_Proteobacteria* | 52 | 186 | 97 | 6.82 | 1.47 | 0.22 |
| Network Suc: 94 nodes and 87 edges | | | | | | | | |
| | *p_Patescibacteria* | *p_Patescibacteria* | 4 | 4 | 1 | 0.14 | 1.15 | 8.37 |
| | *p_Verrucomicrobia* | *p_Verrucomicrobia* | 7 | 7 | 3 | 0.48 | 3.45 | 7.18 |
| | *p_Chloroflexi* | *p_Chloroflexi* | 19 | 19 | 18 | 3.91 | 20.69 | 5.29 |
| | *p_Latescibacteria* | *p_Actinobacteria* | 1 | 14 | 1 | 0.32 | 1.15 | 3.59 |
| | *p_Planctomycetes* | *p_Verrucomicrobia* | 2 | 7 | 1 | 0.32 | 1.15 | 3.59 |
| | *p_Acidobacteria* | *p_Bacteroidetes* | 3 | 15 | 3 | 1.03 | 3.45 | 3.35 |
| | *p_Bacteroidetes* | *p_Bacteroidetes* | 15 | 15 | 6 | 2.40 | 6.90 | 2.87 |
| | *p_Acidobacteria* | *p_Actinobacteria* | 3 | 14 | 2 | 0.96 | 2.30 | 2.39 |
| | *p_Acidobacteria* | *p_Proteobacteria* | 3 | 23 | 3 | 1.58 | 3.45 | 2.18 |
| | *p_Bacteroidetes* | *p_Proteobacteria* | 15 | 23 | 14 | 7.89 | 16.09 | 2.04 |
| | *p_Chloroflexi* | *p_Gemmatimonadetes* | 19 | 6 | 1 | 2.61 | 1.15 | 0.44 |
| | *p_Bacteroidetes* | *p_Chloroflexi* | 15 | 19 | 2 | 6.52 | 2.30 | 0.35 |
| | *p_Proteobacteria* | *p_Chloroflexi* | 23 | 19 | 1 | 10.00 | 1.15 | 0.11 |
| Network Mat: 213 nodes and 206 edges | | | | | | | | |
| | *p_Others* | *p_Patescibacteria* | 10 | 7 | 5 | 0.31 | 2.43 | 7.83 |
| | *p_Verrucomicrobia* | *p_Verrucomicrobia* | 12 | 12 | 3 | 0.29 | 1.46 | 4.98 |
| | *p_Chloroflexi* | *p_Chloroflexi* | 31 | 31 | 14 | 2.06 | 6.80 | 3.30 |
| | *p_Bacteroidetes* | *p_Patescibacteria* | 32 | 7 | 6 | 0.99 | 2.91 | 2.94 |
| | *p_Actinobacteria* | *p_Chloroflexi* | 31 | 31 | 4 | 4.26 | 1.94 | 0.46 |
| | *p_Bacteroidetes* | *p_Verrucomicrobia* | 32 | 12 | 1 | 1.70 | 0.49 | 0.29 |
| Network Old: 363 nodes and 1027 edges | | | | | | | | |
| | *p_Chloroflexi* | *p_Chloroflexi* | 63 | 63 | 118 | 2.97 | 11.49 | 3.87 |
| | *p_Bacteroidetes* | *p_Chloroflexi* | 48 | 63 | 18 | 4.60 | 1.75 | 0.38 |
| | *p_Gemmatimonadetes* | *p_Proteobacteria* | 9 | 127 | 6 | 1.74 | 0.58 | 0.34 |
| | *p_Proteobacteria* | *p_Others* | 127 | 8 | 5 | 1.55 | 0.49 | 0.31 |
| | *p_Actinobacteria* | *p_Bacteroidetes* | 64 | 48 | 13 | 4.68 | 1.27 | 0.27 |
